# Modeling vehicle collision risk for the jungle cat in the Hyrcanian forests of Iran: A guide for vehicle collision prevention

**DOI:** 10.1101/2024.11.27.625583

**Authors:** Abbas Ashoori, Anooshe Kafash, Koros Rabiei, Mojtaba Hosseini, Shapour Abdi, Masoud Yousefi

**Author notes:** Corresponding author: Masoud Yousefi, Department of Animal Science, School of Biology, Damghan University, Damghan, Iran.

## Abstract

Wildlife-vehicle collisions are an important wildlife conservation challenge, especially for carnivores. As in other countries, vehicle collisions pose a major threat to carnivores in Iran. The jungle cat (*Felis chaus*) is a small carnivore species facing multiple threats, including habitat destruction, land use changes, and particularly vehicle collisions. We collected data on jungle cat collisions to model jungle cat-vehicle collision risk in the Hyrcanian forests of northern Iran. We also modeled the jungle cat vehicle collision risk in the study area but by creating 1 km and 5 km buffers around the roads and identified high vehicle collision risk areas within the 1 km and 5 km butters. We used the Maxent model to identify most important predictors of collision risk. Our models showed that areas in west of Golestan province, east of Mazandaran province, and in central parts of Gilan province faced highest vehicle collision risk for the jungle cat in the Hyrcanian forests. Human footprint and slope were the most important predictors of the jungle cat vehicle collision, with 48.3% and 17.2% contribution and with positive and negative correlation respectively. Results of variable importance were similar when modeling area was limited to a 5 km buffer zone around the roads. But when modeling area was limited to a 1 km buffer zone around the roads, slope became insignificant. Collisions more likely occur where vegetation grows immediately adjacent to roads, so clearing a roadside strip would reduce potential collisions. We recommend that high collision-risk areas we identified for jungle cat in the Hyrcanian forests be a focus for future monitoring and conservation planning.

## 1 Introduction

Human development including roads pose serious threat to biodiversity worldwide (Forman et al., 2003; Coffin, 2007; Laurance et al., 2009; Gunson et al., 2011; Laurance et al., 2014; Ward et al., 2015; Baxter-Gilbert et al., 2015; Bennett, 2017; Medrano-Vizcaíno et al., 2022). Mobile species, including carnivores, are particularly vulnerable to road development in human-modified landscapes (Packer et al., 2013; Ripple et al., 2014). Roads may substantially diminish carnivore populations and distributions by increasing mortality as a result of vehicle collision, creating barriers to movement, changing animal behaviours and habitat use patterns, altering the physical and chemical variables, increasing ambient noise, and decreasing the quantity and quality of habitats (Trombulak and Frissell, 2000; Forman et al., 2003; Jaeger and Fahring, 2004; Laurance et al., 2014). Road density also can alter animal movements, fragment populations, and increase human access, and thereby disturb wildlife (Iuell, 2003). Because of these many threats, conservation biologists are evaluating the impacts of roads on species and developing strategies to reduce their negative impacts on biodiversity worldwide (Forman et al., 2003; Ward et al., 2015; Bennett, 2017). For example, recent recognition of the substantial impacts of roads on large mammal populations has prompted many efforts to design and realign roads to reduce road mortality (Gunson et al., 2011; Neumann et al., 2012; Wright et al., 2020; Vilela et al., 2020; Silva et al., 2021). Most efforts to study and reduce negative impacts of roads on biodiversity have occurred in developed countries (Langbein et al., 2011; Gunson et al., 2011; Neumann et al., 2012; Marcantonio et al., 2013; Laurance et al., 2014; Ward et al., 2015; Fabrizio et al., 2019; Loraamm et al., 2019; Wright et al., 2020; Grilo et al. 2020). Road collisions are well documented in these countries (Langbein et al., 2011; Wilkins et al. 2019). For instance, 51,522 animal collisions were recorded in Texas, from 2010 to 2016, these resulted in 254 human fatalities and 6,914 injuries, in addition to animal loss (Wilkins et al. 2019). Species Distribution Models (SDMs) are being used to identify areas showing high road collisions probability and what are the environmental predictors of road collisions within the defied geographic area (Fabrizio et al., 2019; Wright et al., 2020). Wright *et al*. (2020) used SDMs and identified areas showing a high probability of European hedgehog (*Erinaceous europaeus*) roadkill occurrence throughout the entire British road network. In another study, Fabrizio et al. (2019) applied SDMs to determine roadkill risk areas for the Eurasian badger (*Meles meles*) in the Abruzzo region (Central Italy). In contrast, the impacts of roads on biodiversity remained relatively unknown in many developing countries compared to developed countries (Parchizadeh et al., 2018). Previous studies have shown that the development of road networks in Iran has been a serious conservation issue for many wildlife species (Moqanaki and Cushman, 2017; Mohammadi et al., 2018; Parchizadeh et al., 2018; Naderi et al., 2018). For example, Naderi et al. (2018) documented and modeled anthropogenic mortality sources in the Persian leopard (*Panthera pardus*) in Iran and found that road mortality was the second most frequent cause of unnatural mortality. They also showed that the species mortality risk is higher in northern Iran than elsewhere in the country (Naderi et al., 2018). Road mortality also has been identified as a risk to the jungle cat (*Felis chaus*; Sanei et al., 2016).

Northern Iran is largely covered by relict deciduous. Hyrcanian forests which comprise a continuous 800-kilometer, with an area of over 1.8 million hectares (Akhani et al., 2010; Sagheb Talebi et al., 2014; Moradi et al., 2019; Yousefi et al., 2023). This area is rapidly changing due to urban and agricultural development, and resulting human population increase (Mahmoudi et al., 2016; Meinecke et al., 2018). As a result, it supports species that are among the most vulnerable in Iran (Meinecke et al., 2018; Ashoori et al., 2018). Hyrcanian forests support high biological diversity and endemism (Akhani et al., 2010; Mahmoudu et al., 2016; Soofi et al., 2019; Kafash et al., 2021, 2022; Jouladeh-Roudbar et al., 2020; Yousefi et al., 2022) because it served as a refugia for species during ice ages (reviewed in Yousefi et al., 2023). Hyrcanian forests are home to many endangered and ecologically important mammals including Persian leopard, Eurasian lynx (*Lynx lynx*), jungle cat and probably the wild cat (*Felis lybica*) (Karami et al., 2016). Climate and land use changes, grazing, illegal hunting, and road mortality are important anthropogenic causes of biodiversity loss in this unique ecosystem (Sagheb Talebi et al., 2014; Meinecke et al., 2018; Naderi et al., 2018; Yousefi et al., 2019; Soofi et al., 2019). Jungle cat mortality rates in the area were reported to be higher than other parts of the country and therefore warranted monitoring.

Our goal was to develop a vehicle collision risk model for the jungle cat in the Hyrcanian forests, to identify the most influential environmental factors contributing to vehicle collision risk, and identify high vehicle collision risk areas within and outside protected areas. We hypothesized that vehicle collision risk is higher in areas with higher human development and road density. We also hypothesized that jungle cat vehicle collision would be higher in areas of dense roadside vegetation cover because of lower sight distance by both driver and cats in such areas.

## 2 Material and methods

### 2.1 Study area

The study areas consisted of Hyrcanian forests in three Iranian provinces: Gilan, Mazandaran and Golestan (Figure 1). The area extends along the Caspian Sea from the boundary of Iran and Azerbijan Republic on the southwest side to Golestan National Park on the southeast side (Figure 1). Mean annual precipitation ranges mainly between 530 and 1350 mm but reaches up to 2000 mm in the west (Sagheb-Talebi et al., 2014). Thus, the area is substantially wetter than most of the rest of Iran, where precipitation averages 250 mm per year. The study area has a mild subtropical humid climate that is favorable for agriculture and includes a center of food production for Iran (Kosarev, 2005). The area as a whole is largely rural, composed of small towns and low-density farmlands and grazing lands.

**FIGURE 1.**
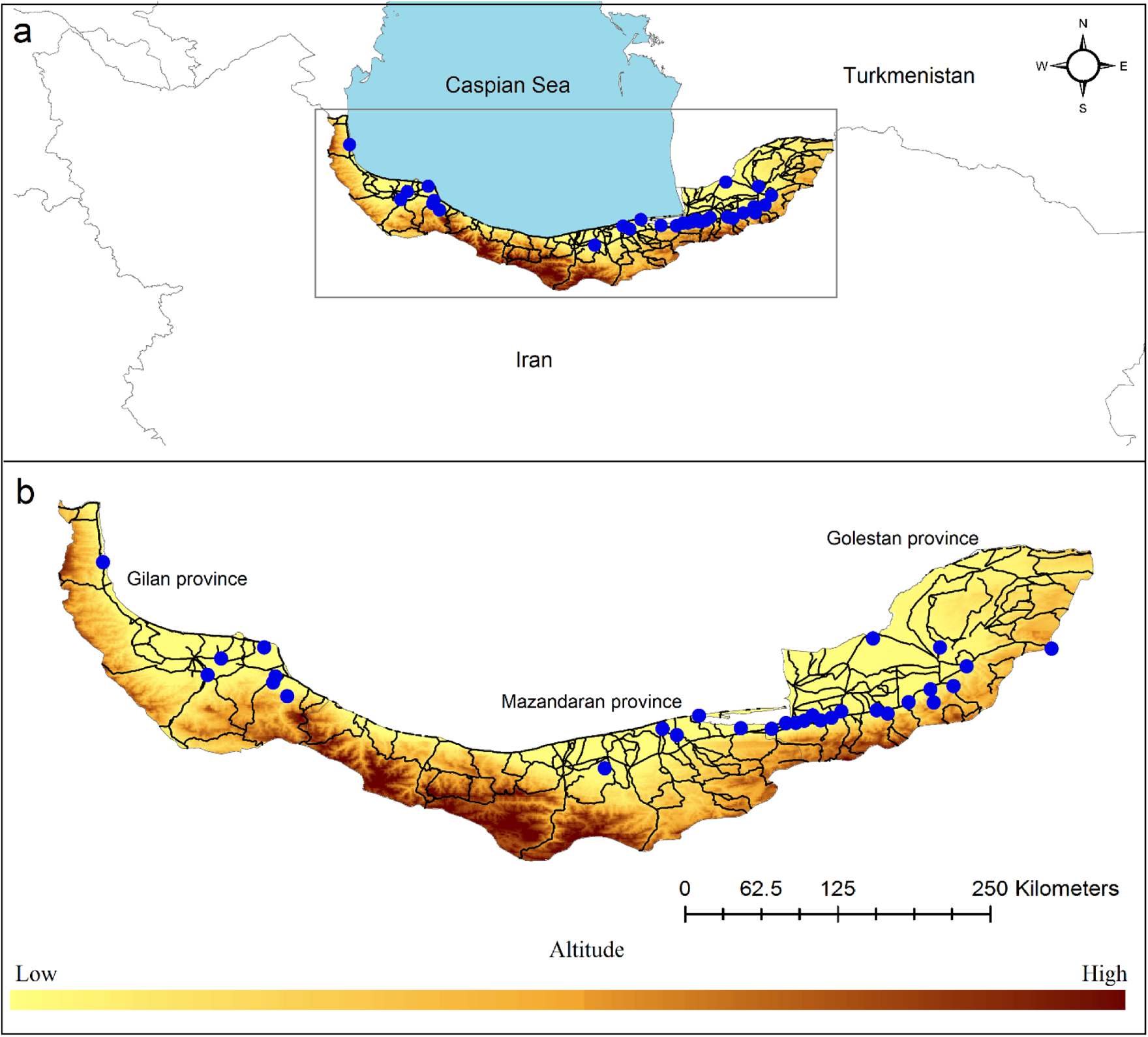
Location of study area in Iran (a). A topographic overview of study area, altitude, roads and jungle cat vehicle collision locations (b). The study areas consisted of Hyrcanian forests in three Iranian provinces: Gilan, Mazandaran and Golestan.

### 2.2 Study species

The jungle cat occurs in North Africa and is widespread in Asia from the Middle East, Southwest Asia, Central and South Asia to Southeast Asia, reaching Indochina and possibly the Malayan Peninsula (Nowell and Jackson, 1996; Abu-Baker et al. 2003, Duckworth et al., 2005). The species has been reported to occur throughout most of Iran except in deserts (Karami et al., 2016; Sanei et al., 2016). The species occurs at elevations from -28 m to 4,178 m asl., the widest elevational distribution of the eight living felid species of Iran (Karami et al., 2016; Sanei et al., 2016). Despite this wide distribution, the species is among the least known cat species in Iran (Karami et al., 2016, Sanei et al., 2016;). Only a single study has been published on the distribution, characteristics, and conservation of the jungle cat in Iran (Sanei et al., 2016). The population of the species in Iran has been reported to have significantly decreased over the past years (Ziaie, 2008). It is cited in the Appendix II of CITES and is legally protected in Iran (Karami et al., 2016).

### 2.3 The jungle cat vehicle collision data

Vehicle collisions records were collected opportunistically from March 2016 to April 2019 in north of Iran (Figure 1). We registered coordinates of 30 collision site when we could confidently confirm the identification of the species using Global Positioning Systems (GPS) (available at https://doi.org/10.5281/zenodo.10372265). We then checked the occurrence data of collision sites for spatial autocorrelation and applied the global Moran’s test to evaluate the structural pattern of the occurrence data.

### 2.4 Environmental Data

We used 11 climatic, topographic, anthropogenic, and Normalized Difference Vegetation Index (NDVI), and variables to quantify environmental characteristics of collision sites and identify the most important predictor of collision risk. Climatic variables were downloaded from WorldClim database at 30-seconds spatial resolution (Fick and Hijmans, 2017) including: isothermality (Bio3), seasonal temperature change (Bio4), mean temperature of wettest season (Bio8), mean temperature of driest season (Bio9), seasonal precipitation change (Bio15), precipitation of warmest season (Bio18), and precipitation of coldest season (Bio19). We used NDVI as an indicator of primarily productivity and vegetation cover within the study area. We considered the human footprint index (Venter et al., 2016a) as a measure of anthropogenic impact. This index is based on the extent of built environments, including population density, electrical infrastructure, croplands, pasture lands, roads, railways, and navigable waterways (Venter et al., 2016b). Slope and topographic heterogeneity were obtained from the Shuttle Radar Topography Mission (SRTM) elevation model (Jarvis et al., 2008). We calculated variance inflation factor (VIF; Quinn and Keough, 2002) for the variables using ‘usdm’ package (Naimi, 2015) in R 3.6.0 environment (R Core Team, 2016). The results showed that collinearity was low (VIF<10) among variables (Bio3 = 3.399, Bio4 = 6.38, Bio8 = 7.917, Bio9 = 8.127, Bio15 = 1.833, Bio18 = 6.42, Bio19 = 3.739, slope = 5.882, topographic heterogeneity = 3.502, NDVI = 1.885, human footprint = 1.756).

### 2.5 Vehicle collision risk modeling

We used Maxent 3.4.1 (Phillips et al., 2019) to predict areas with high vehicle collision risk in the study area (Phillips et al., 2006; Phillips and Dudík, 2008). We ran Maxent with maximum iterations of 500, a convergence threshold of 0.0001 and 1000 background points (available at https://doi.org/10.5281/zenodo.10372265). We used the cross-validation method in Maxent. Maxent determines the importance of each environmental variable as a value between 0 to 100 percent (Phillips et al., 2006). We built two models to identify the most influential predictor of vehicle collision risk for the jungle cat. The first model was developed using the eleven variables (Bio3, Bio4, Bio8, Bio9, Bio15, Bio18, Bio19, slope, topographic heterogeneity, NDVI, human footprint) and then second model was run only by including those variables that contributed more than 1% to the first model (Yousefi et al., 2018). We also modeled the jungle cat vehicle collision risk in the study area but by creating 1 km and 5 km buffers around the roads and identified high vehicle collision risk areas within the 1 km and 5 km butters. Distribution points were randomly split into 10 folds containing equal number of occurrences, and training models were created by eliminating each fold in turn (Merow et al., 2013). The performance of the vehicle collision risk models was assessed using the area under the curve (AUC) metric of the receiving operator characteristic (ROC) curve (Phillips et al., 2006). AUC is commonly used as a measure of model performance in ecological studies (Phillips and Dudík, 2008; Heumann et al., 2011; Kafash et al., 2018). AUC ranges from 0 to 1 an AUC value of 0.5 indicates that the performance of the model is not better than random, while values closer to 1 indicate better model performance (Swets, 1988).

### 2.6 Representation of high vehicle collision risk areas within protected areas

To estimate locations with high vehicle collision risk for the jungle cat within protected areas, we first converted the continuous vehicle collision risk map to a binary map using the maximum test sensitivity plus specificity threshold (cut of value 0.21) (Phillips et al., 2006; Phillips et al., 2019). Then we overlaid the binary map with protected areas of Iran (Darvishsefat, 2008) and calculated vehicle collision risk area within protected areas network. The most recent shapefile of Iran’s protected areas was obtained from Department of Environment of Iran.

## 3 Results

### 3.1 Vehicle collision risk model

We recorded 30 jungle cat mortality incidents from the three northern Iran provinces. Based on the AUC metric of the ROC curve, the overall predictability of the jungle cat vehicle collisions model was high (AUC = 0.906 ± 0.014). The Maxent model showed that the highest risk of vehicle collisions for the jungle cat in the study area occurred within western and central Golestan, eastern Mazandaran and central Gilan (Figure 2).

**FIGURE 2.**
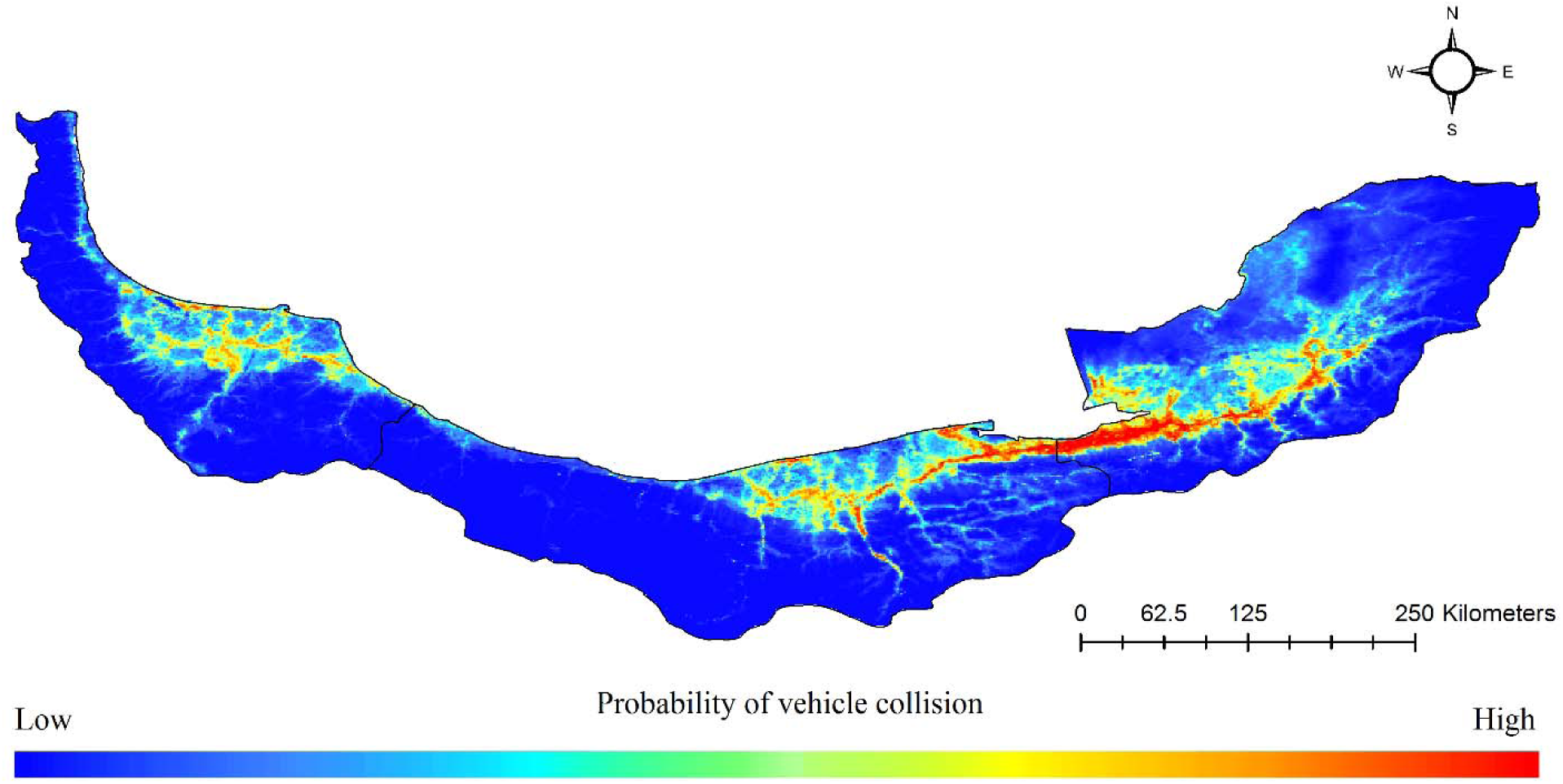
Relative probability of the jungle cat vehicle collisions across the study area. The collision risk model is developed based on 30 jungle cat vehicle collision locations and Maxent model by considering following environmental variables: human footprint, slope, topographic heterogeneity, NDVI, distance to wetlands, distance to rivers, isothermality (Bio3), seasonal temperature change (Bio4), mean temperature of wettest season (Bio8), mean temperature of driest season (Bio9), seasonal precipitation change (Bio15), precipitation of warmest season (Bio18), precipitation of coldest season (Bio19) at 30-seconds spatial resolution.

### 3.2 Road based vehicle collision risk models

We also predicted the jungle cat vehicle collision risk areas by creating 1 km and 5 km buffers around the roads. We found that roads crossing western and central Golestan, eastern Mazandaran and central Gilan are associated with highest vehicle collision risk (Figure 3). Based on the AUC metric of the ROC curve, the overall predictability of the jungle cat vehicle collisions models at 5 km buffer (AUC = 0.841 ± 0.027) and 1 km buffer (AUC = 0.883 ± 0.019) was high.

**FIGURE 3.**
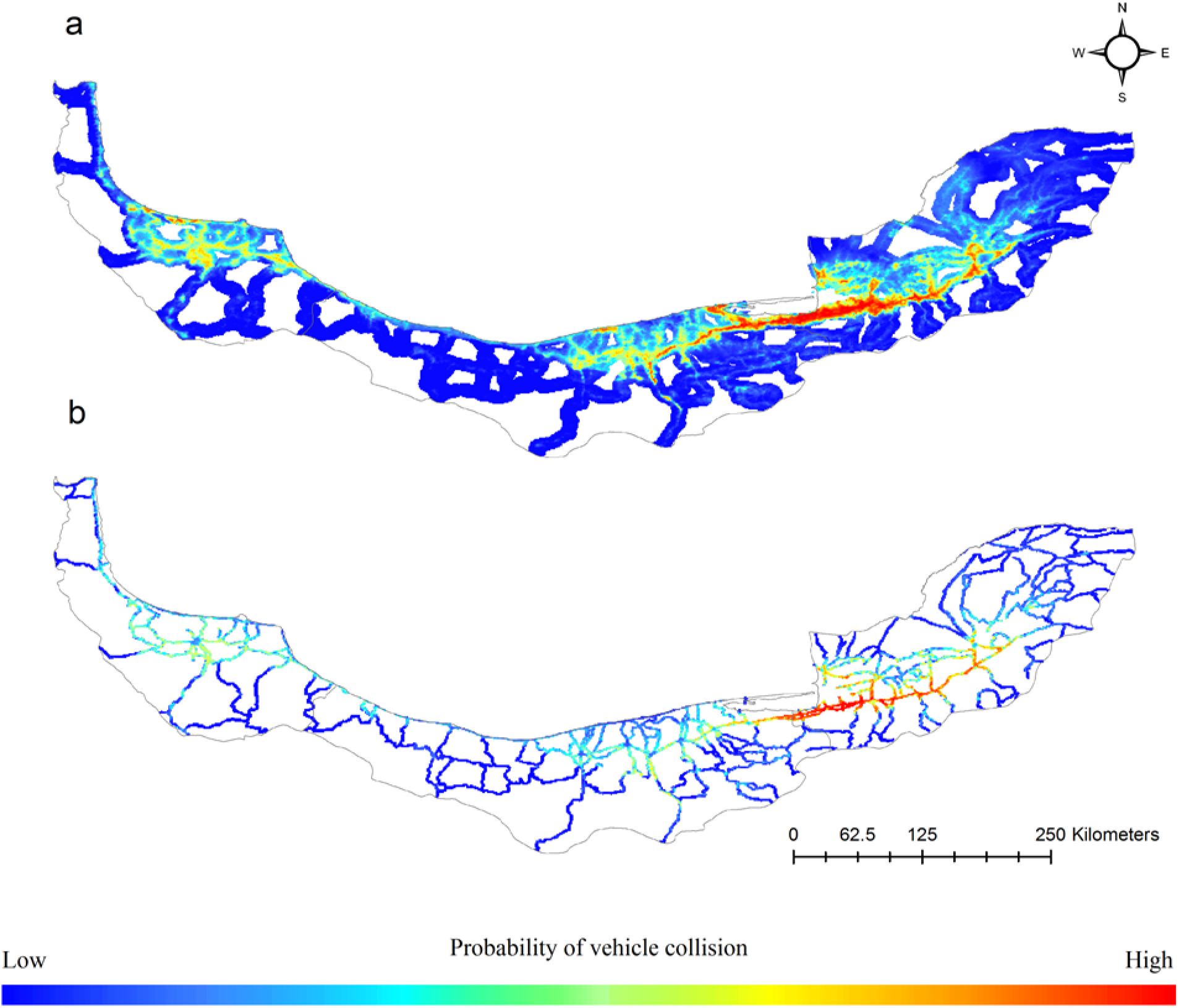
Relative probability of the jungle cat vehicle collisions based on Maxent model based on the 5 km (a) and 1 km (b) buffers around the roads. The models were created by considering following environmental variables: human footprint, slope, topographic heterogeneity, NDVI, distance to wetlands, distance to rivers, isothermality (Bio3), seasonal temperature change (Bio4), mean temperature of wettest season (Bio8), mean temperature of driest season (Bio9), seasonal precipitation change (Bio15), precipitation of warmest season (Bio18), precipitation of coldest season (Bio19) at 30- seconds spatial resolution.

### 3.3 Variable contribution

Human footprint and slope were the most important predictor of the jungle cat vehicle collision, with 48.3% and 17.2% contributions respectively (Table 1). Human footprint and jungle cat vehicle collision risk were positively correlated (Figure 4). Slope and jungle cat vehicle collision risk were negatively correlated, with lower incidence of collisions in steeper areas. The probability of jungle cat vehicle collisions also increased with an increase in NDVI values while the probability decreased with an increase in distance from wetlands and rivers. Results of variable importance were similar when modeling area was limited to a 5 km buffer zone around the roads. But when modeling area was limited to a 1 km buffer zone around the roads, slope which was the second most important predictor became insignificant however, human footprint was still the most important predictor of the jungle cat vehicle collision.

**FIGURE 4.**
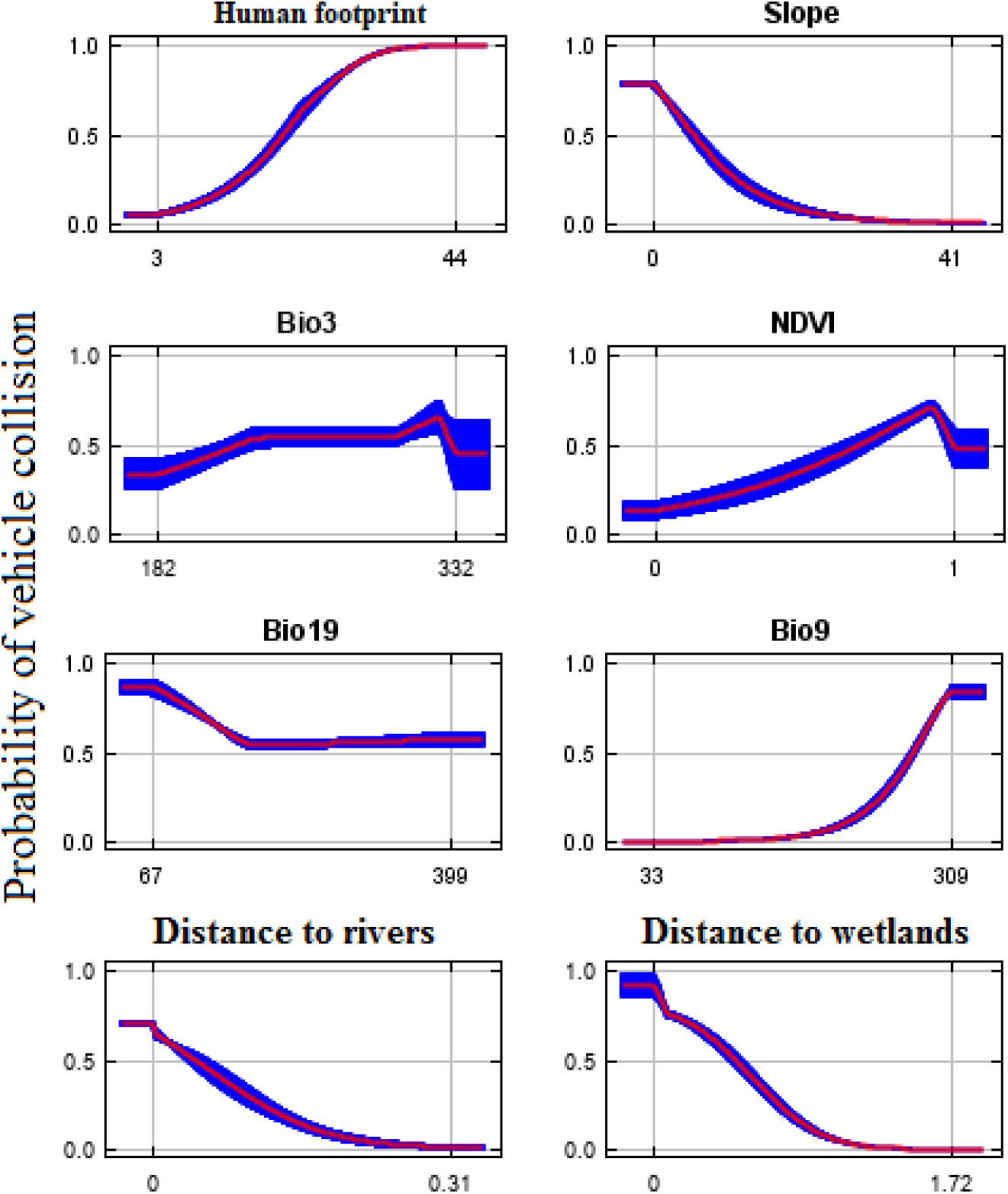
Response curves for those variables in the jungle cat vehicle collisions model having more than 1.0% contribution. Key to symbols: Bio3 = isothermality, Bio9 = Mean temperature of driest season, Bio19 = precipitation of coldest season.

**TABLE 1.**
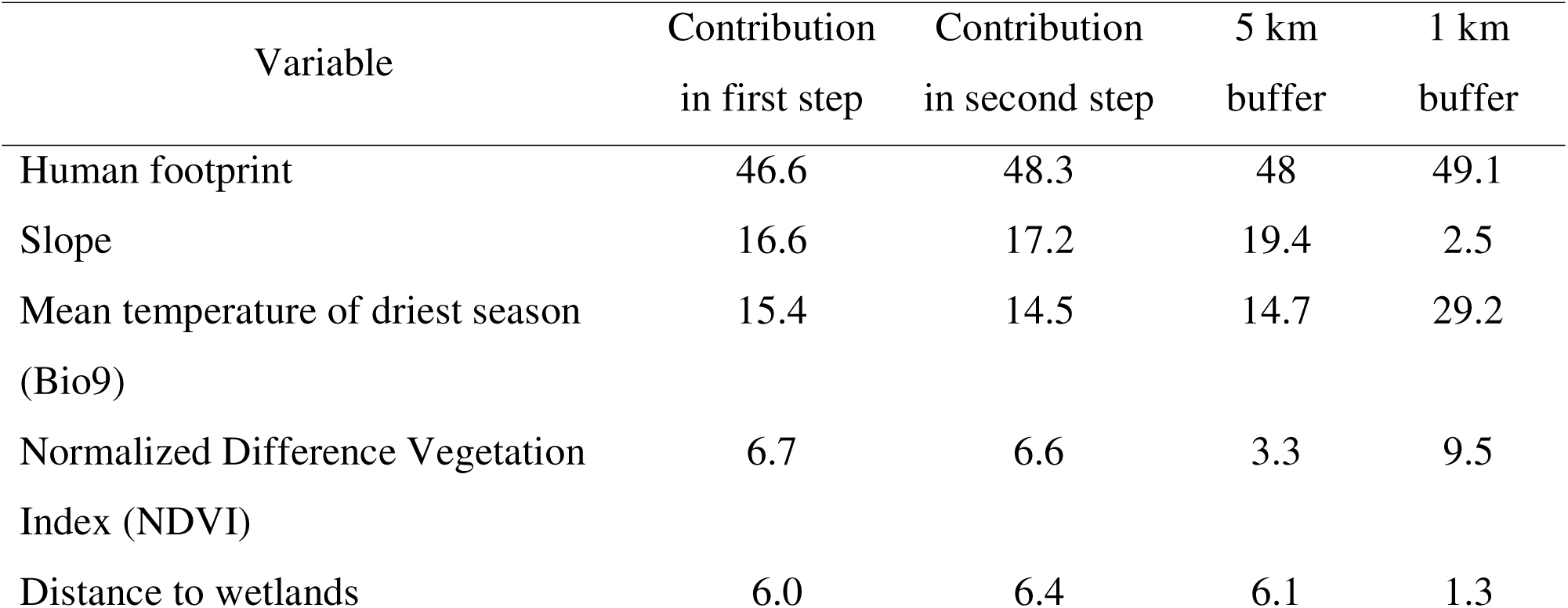
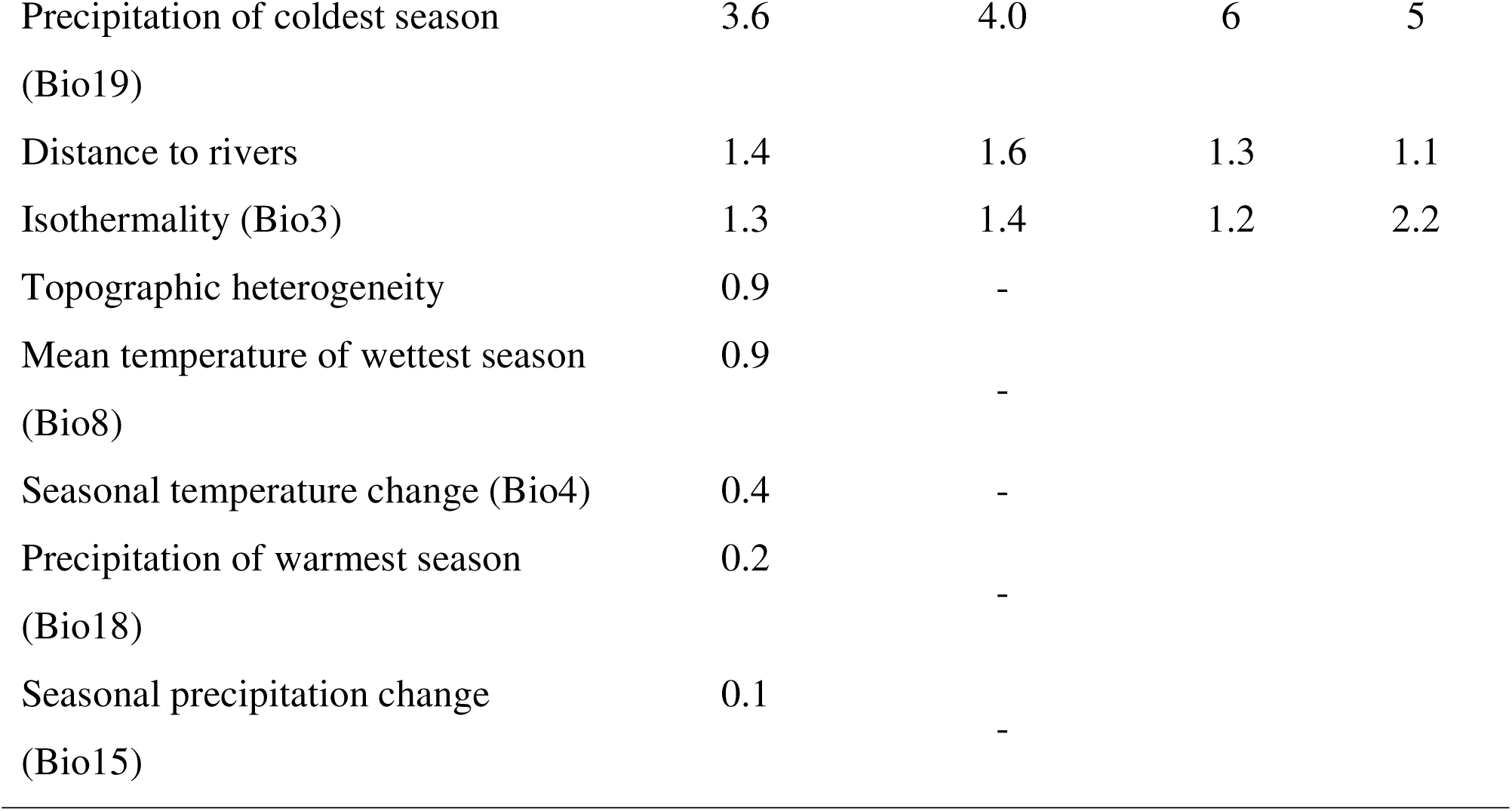
Contributions of environmental variables to the vehicle collision risk model for the jungle cat in the Hyrcanian forests, northern Iranian provinces.

| Variable | Contribution<br>in first step | Contribution<br>in second step | 5 km<br>buffer | 1 km<br>buffer |
| --- | --- | --- | --- | --- |
| Human footprint | 46.6 | 48.3 | 48 | 49.1 |
| Slope | 16.6 | 17.2 | 19.4 | 2.5 |
| Mean temperature of driest season<br>(Bio9) | 15.4 | 14.5 | 14.7 | 29.2 |
| Normalized Difference Vegetation<br>Index (NDVI) | 6.7 | 6.6 | 3.3 | 9.5 |
| Distance to wetlands | 6.0 | 6.4 | 6.1 | 1.3 |
| Precipitation of coldest season<br>(Bio19) | 3.6 | 4.0 | 6 | 5 |
| Distance to rivers | 1.4 | 1.6 | 1.3 | 1.1 |
| Isothermality (Bio3) | 1.3 | 1.4 | 1.2 | 2.2 |
| Topographic heterogeneity | 0.9 | - |  |  |
| Mean temperature of wettest season<br>(Bio8) | 0.9 | - |  |  |
| Seasonal temperature change (Bio4) | 0.4 | - |  |  |
| Precipitation of warmest season<br>(Bio18) | 0.2 | - |  |  |
| Seasonal precipitation change<br>(Bio15) | 0.1 | - |  |  |

### 3.4 Protected areas coverage

We calculated the extent to which areas of high vehicle collision risk for the jungle cat occurred within protected areas. Results showed that 13,878 km^2^ of the study areas have high vehicle collision risk for the species and only 213 km^2^ (1.5 percent) of the high risk areas were located within protected areas (Figure 5).

**FIGURE 5.**
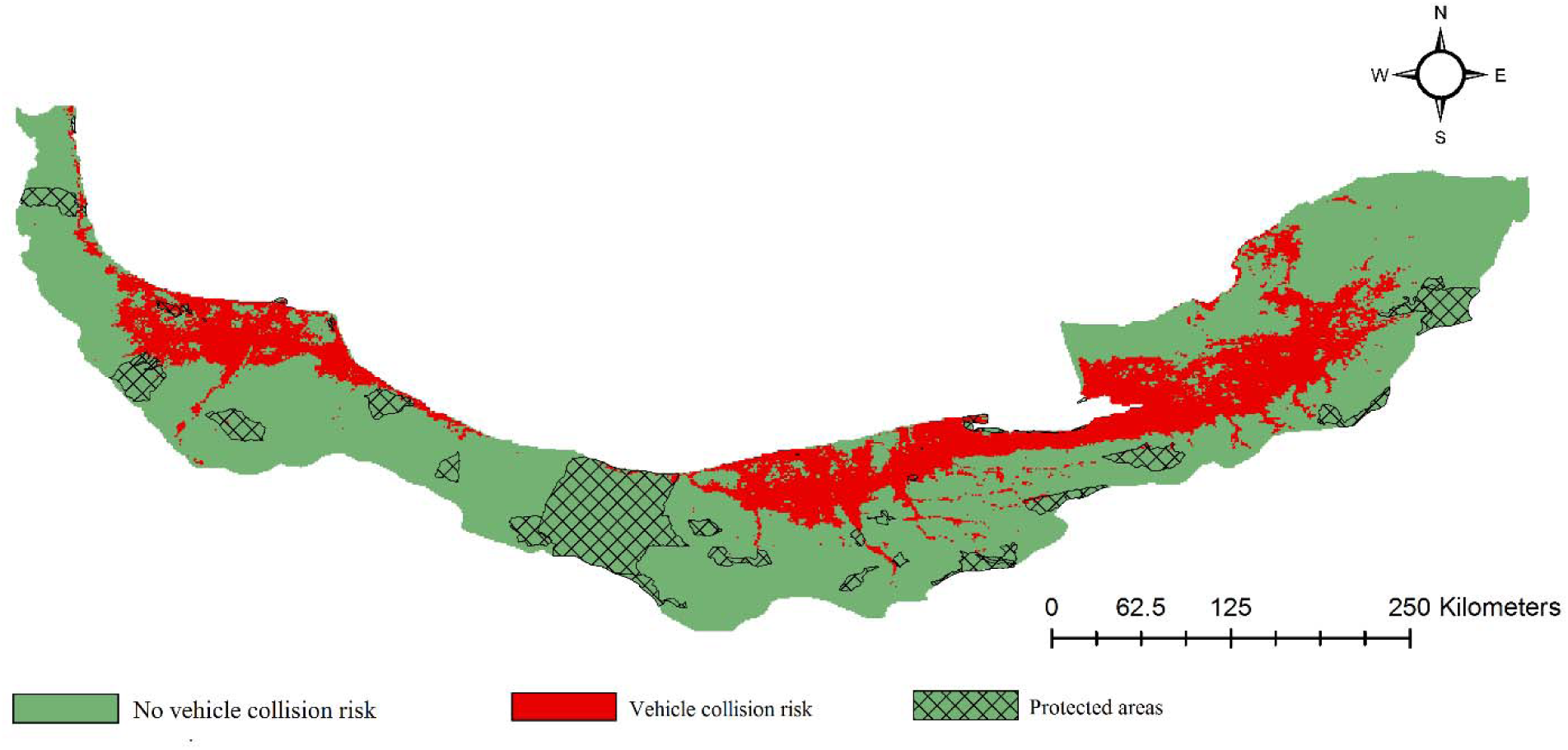
Relative risk of vehicles for the jungle cat in relation to protected areas.

## 4 Discussion

Road development is a serious threat to carnivore species (Meza-Jova et al., 2019 because of their large home ranges and need to maintain population. Therefore, understanding the measures of this phenomenon is vital to find spatial distribution o roadkill hotspots and to mitigate its effect on carnivore population (Miranda et al., 2020). Our quantification of environmental characteristics associated with jungle cat vehicle collisions risk found that human footprint and slope were the main predictors of vehicle collision risk.

Human footprint variable was the most important determinant of the jungle cat vehicle collision risk and, as expected, when extent of the human footprint increased the probability of vehicle collision also increased. The relationship between human footprint and jungle cat mortality is somewhat surprising, in that areas of highest human footprint would be expected to be too degraded to provide suitable or high value habitat for the species. The species avoids areas with the highest human density especially highly urbanized areas, but it utilizes the outskirts of urban areas, and readily penetrates smaller cities and villages which are the most typical form of human development in the study area. In fact, the cat appears to favor small urban areas and villages because of the augmented availability of prey such as domestic chickens (*Gallus gallus*). This use pattern demonstrates the species’ tolerance of some degree of habitat modification (Duckworth et al., 2005), as shown by records of breeding within villages (A. Ashoori, pers. obs.).

Slope was the second most important predictor of the jungle cat vehicle collision risk in Hyrcanian forests. The relationship showing that mortalities primarily occur on low slopes could be interpreted to suggest that roads are more likely to occur on flatter lands, rather than reflecting the response of the species to slope (Karami et al., 2016). The complete absence of road kill reports from mountainous areas, as well as a previous study (Mousavi et al., 2018), both indicate that the jungle cat avoids steep slopes and higher elevation areas (Karami et al., 2016). Slope was also the second most important determinant of the species vehicle collision risk when modeling areas was limited to a 5 km buffer zone around roads but it did not come out to be important when modeling areas was limited to a 1 km buffer zone around the roads.

Plant cover plays a key role to the life of jungle cat, which favoured woodlands and shrublands more than other land cover types (Sanei et al., 2016). Our results demonstrate that plant cover adjacent to roads could increases the frequency of roadkill. Therefore, keeping lower plant cover around roads could probably reduce the rate of roadkill. Similarly, the rate of roadkill of European hedgehog (*Erinaceous europaeus*) increased with the extent of grassland cover (Wright et al., 2020). Therefore, conserving open forests along roadsides, which could provide cover favored by the jungle cat but also increase visibility among cats and vehicles is recommended (Duckworth et al., 2005). Removal of plant cover adjacent to roads, however, may also increase the barrier effect of roads to jungle cat movement. More research is needed to understand how roads and vegetation clearing along roads may fragment habitats and affect road mortality.

We acknowledge certain limitations in interpreting and applying our study results. Although it may be obvious, it is important to stress that vehicle mortality records represent the intersection of the distributions of the jungle cat and road network. The data do not represent an unbiased estimate of the range or abundance of the jungle cat in any particular geographic region, but only the range and abundance of vehicle collision mortality within sampled areas. Therefore, caution is required in interpreting relationships between road mortalities and certain variables, as relationships may more reflect conditions where roads occur rather than where jungle cats occur. For example, the strong relationship between human footprint and cat mortalities may not reveal anything about the species’ population response to development. Rather, the result may just indicate that the presence of more humans, roads, and vehicles result in higher mortality regardless of the density of species.

Our results suggest several worthwhile areas for further research. Determining the density of roads in suitable habitat could be helpful in further determining what proportion the species range is subject to roadkill. Quantifying the number of kills/road mile/year could also improve the characterization of potential population impacts. Developing a systematic reporting system for road mortalities by enlisting highway maintenance workers and interested community members, would provide a better indication of the frequency of roadkills, areas of the species range with higher and lower roadkill rates, and roadside conditions associated with vehicle mortalities. Further information derived from roadkill hotspots should be helpful in mitigating when construction of new roads (such as locating road alignments away from riparian and wetland habitats) but offers fewer options for existing roads (i.e., warning signs, installing lighting, vegetation clearing, enhancement of existing undercrossings to facilitate movements). Future studies are needed to more finely quantify conditions at the immediate site of impact (i.e., number of road lanes, traffic volume, and multi-lane roads) as well as jungle cat density in areas where roadkills occur. Studies of the effectiveness of measures to reduce mortalities are also needed. For example, vegetation clearing and installation of lighting at areas of high roadkill risk may reduce mortalities, but it also may prevent or reduce jungle cats from using these areas, potentially resulting in population fragmentation. Our mapping of mortalities at existing roads may partially reflect conditions created by impacts of previous roadkill. As a result, roadkill records could have shifted from high-traffic to low-traffic segments, obscuring the full extent of roadkill impacts on the species (Zimmermann et al., 2017).

Results of our evaluation of the overlap of collision occurrence and risk and protected areas deserve careful interpretation. The location of all collision incidents and most areas of high collision risk outside of protected areas demonstrates that collision risk is not high in these areas, but we only have limited knowledge that suggests that these areas, based on their location in more montane topography, may support lower densities of jungle cat than unprotected lowlands. If protected areas do not support high jungle cat densities, additional land protection may be warranted to protect the species in areas where it is now at risk from vehicle collisions.

## 5 Conclusion

We showed that areas with a high human footprint and low slope around road network in Hyrcanian forests have high vehicle collision risk for the jungle cat. Although 98.5 percent of high vehicle collision risk areas are outside protected areas, the presence of many protected lands in areas of high slope (Darvishsefat, 2008) makes it uncertain to what degree these protected areas support jungle cat populations. Using roadside fences to direct jungle cat movements to areas below roads (under bridges or through culvets) could be effective in reducing road crossings and resulting vehicle collisions. We also recommended experimental treatments to reduce vegetation cover in zone from ground level to 2 m by tree thinning or removal of shrub and herbaceous vegetation beside roads near river crossings and wetlands, where jungle cats are most likely to occur. Partial overhead tree cover should be retained in these areas because the species appears to prefer such areas and the presence of tree cover also will discourage the growth of low vegetation and thereby retain desirable conditions and reduce the frequency of vegetation treatment. As roads appear to be a threat for the jungle cat, we recommend quantifying vehicle collision risk across the species distribution range in Iran to be able to apply measures to reduce road mortality.

## Acknowledgements

We are grateful for the help of the Environment Department of Gilan, Mazandaran and Golestan Provinces. We thank Mohamad Sadegh Khosravi, Yaghoub Rakhshbhar and Fardin Naziri for supplying several of the vehicle collision locations. We thank Iran National Science Foundation for financial support (Project number: 4004303). We also thank Daniel A. Airola for his careful reading of our manuscript and help in improving it.

## Author contributions

**Abbas Ashoori**: Writing - Original Draft (equal); investigation (equal); formal analysis (equal). **Anooshe Kafash**: Writing - Original Draft (equal); investigation (equal); formal analysis (equal); software (lead); methodology (lead). **Koros Rabiei**: investigation (equal). **Mojtaba Hosseini**: investigation (equal). **Shapour Abdi**: investigation (equal). **Masoud Yousefi**: Conceptualization (lead); formal analysis (lead); investigation (equal); methodology (equal); Visualization (lead); writing – original draft (lead); writing – review and editing (lead).

## Data availability statement

The data that support the findings of this study are presented at Zenodo: https://doi.org/10.5281/zenodo.10372265.

